# Time-series foundation modeling enables accurate lake ecosystem forecasting

**DOI:** 10.64898/2026.09.13.751332

**Authors:** Asuka Matsushita, Hideyuki Doi

## Abstract

Accurate ecological forecasting is increasingly essential for understanding and managing ecosystem responses to climate variability and anthropogenic pressures; however, prediction remains difficult in lakes because key biological variables, such as phytoplankton biomass, exhibit nonlinear dynamics, observational noise, data sparsity, and non-stationarity. This study aimed to test whether large pre-trained time-series foundation models can improve forecasts of lake phytoplankton dynamics, focusing on chlorophyll-a (Chl.a) and water-quality variables in two ecologically contrasting Japanese lakes, the deep-stratified Lake Biwa and the shallow nutrient-rich Lake Kasumigaura. Using extensive monthly monitoring records spanning up to 30 years, we benchmarked two time-series foundation models—Transformer-based Chronos-T5 and probabilistic Lag-Llama—against 15 statistical (AR, ARIMA, SARIMA, Prophet), machine learning (Random Forest, XGBoost, KNN, SVR), and deep learning (LSTM, CNN, TCN, and SSA-hybrid) approaches. Chronos-T5 achieved superior reproduction accuracy and stability across diverse environmental variables and outperformed all other tested models. This performance stemmed from its ability to capture long-term dependencies and complex temporal patterns through large-scale pre-training. We further identified an optimal training window of approximately 14 years, balancing data sufficiency with regime stability, beyond which model accuracy diminished owing to ecological regime shifts. Our findings highlight the transformative potential of time-series foundation models for ecological forecasting, providing a scalable, data-driven framework that can underpin robust early warning systems and adaptive management strategies for aquatic ecosystems worldwide.

**Significance statement:** Accurate forecasting of lake ecosystems is crucial for water management under climate change, yet predicting phytoplankton dynamics remains challenging due to complex, non-stationary biological responses. Here, we demonstrate that large time-series foundation models significantly outperform traditional statistical, machine learning, and deep learning methods in forecasting chlorophyll-a and water-quality dynamics across contrasting Japanese lakes. Pre-trained on vast temporal datasets, these models successfully capture long-term dependencies and non-linear patterns without domain-specific architecture tuning. Furthermore, we identified an optimal 14-year training window, highlighting how ecological regime shifts constrain model memory. Our findings introduce time-series foundation models as a scalable, high-precision paradigm for ecological forecasting, offering vital tools for global freshwater conservation and early-warning systems.

## Introduction

Predicting ecosystem responses to environmental variability and human pressure is fundamental to modern ecology and natural resource management. As global climate change accelerates environmental alterations, the urgency to anticipate regime shifts, biodiversity loss, and ecosystem service disruptions has intensified ^1–3^. This predictive capacity enables proactive decision-making to mitigate adverse effects and supports adaptive management strategies that enhance the resilience of ecosystems.^4–6^ Moreover, ecological forecasting integrates diverse data sources and modeling approaches to capture complex ecological dynamics across spatial and temporal scales, thereby improving prediction accuracy and relevance. Its applications extend beyond conservation biology to include agriculture, fisheries, and urban planning, highlighting its critical role in sustaining ecosystem functions and services amid rapid environmental change.^1,4,7,8^

Traditional ecological forecasting approaches primarily rely on linear statistical models, such as autoregressive integrated moving average (ARIMA)^1^. These models are computationally efficient but often fail to capture the nonlinear and complex temporal dependencies typical of ecological time series. ^2^ Their assumptions of stationarity and linear relationships limit their effectiveness in dynamic ecosystems influenced by multiple interacting stressors. Machine learning methods, including random forest and XGBoost, represent a significant advancement by enabling the recognition of nonlinear patterns.^9,10^ However, these approaches typically require extensive feature engineering and may struggle to effectively model temporal dependencies and handle data sparsity. While they generally outperform linear models in terms of predictive accuracy, their limited capacity to capture sequential dependencies restricts their suitability for forecasting ecological processes that evolve over time.^11,12^ Deep learning architectures, such as long short-term memory (LSTM) networks and convolutional neural networks (CNN), further enhance forecasting capabilities by modeling sequential dependencies and extracting hierarchical features from time-series data.^13,14^ These models can theoretically capture complex temporal patterns and nonlinear interactions without manual feature engineering. Nevertheless, their performance is often constrained by challenges inherent in ecological datasets, including limited data availability, noise, non-stationarity, irregular sampling intervals, missing data, and high variability. These factors contribute to overfitting and poor generalization when training deep learning models on relatively small ecological datasets.

Recently, the emergence of large-scale time-series foundation models, inspired by advances in natural language processing, has introduced a paradigm shift in time-series forecasting.^15,16^ These models, pre-trained on vast, heterogeneous pre-trained multiple domains, internalize universal temporal patterns, thereby enabling superior zero-shot generalization and data efficiency.^17–19^ By leveraging transfer learning, foundation models can adapt to specific ecological forecasting tasks with limited fine-tuning, thereby reducing their dependency on large labeled datasets. In particular, transformer-based models, such as Chronos-T5, and probabilistic frameworks, such as Lag-Llama, offer promising avenues for ecological forecasting, given their capacity to model long-term dependencies, capture uncertainty, and handle irregular time series.^2,20^ Their attention mechanisms allow for flexible weighting of temporal information, which is crucial for detecting subtle ecological signals in the presence of noise.

Aquatic ecosystems, especially lakes, serve as sentinel systems that reflect broader environmental changes and are vital for freshwater biodiversity and human water supply.^21,22^ Phytoplankton, as primary producers in these ecosystems, play a crucial role in regulating biogeochemical cycles and maintaining water quality, and their population dynamics are central to ecosystem functioning and management efforts.^23,24^ Predicting ecosystem dynamics, including phytoplankton variability, requires considering a comprehensive set of environmental factors, such as chlorophyll concentration, water temperature, nutrient levels (notably nitrogen and phosphorus), and dissolved oxygen (DO). These variables collectively influence the growth rate, species composition, and overall health of the ecosystem. The complex interplay of biological interactions, variable environmental drivers, and data limitations makes prediction challenging.^15,16,25^ Temporal and spatial variability further complicates modeling, as short-term fluctuations may obscure long-term trends that are essential for management. Advances in remote sensing, high-frequency monitoring, and data-driven modeling approaches, including machine learning techniques, have enhanced our ability to capture this multifaceted variability. However, integrating diverse data sources and accounting for nonlinear ecological responses remain significant challenges in this field. In this study, environmental data, including chlorophyll, water temperature, nitrogen and phosphorus nutrients, and dissolved oxygen, were used as example predictors to test the models’ ability to predict ecosystem dynamics. Understanding and forecasting these dynamics is essential for anticipating harmful algal blooms, managing water quality, and conserving aquatic biodiversity under changing climatic and anthropogenic pressures.^15,16^

We aimed to clarify both the advantages and limitations of foundation models in capturing the complex temporal dynamics of phytoplankton populations and evaluate their usefulness for practical ecological forecasting. We propose that foundation models have the potential to overcome traditional data limitations by delivering highly accurate and broadly applicable forecasts, which are crucial for advancing ecological understanding and guiding ecosystem management. This combined approach is essential for converting model predictions into practical guidance that ecosystem managers can use to address and mitigate the effects of environmental changes. In this study, we utilized a 30-year weekly dataset to investigate the application of time-series foundation models for forecasting chlorophyll-a and water quality dynamics in two ecologically contrasting Japanese lakes, Lake Biwa, and Lake Kasumigaura. These lakes differ significantly in terms of trophic status, hydrodynamics, and anthropogenic impacts, providing a rigorous framework for assessing the generalizability of models across diverse ecological conditions. By leveraging extensive long-term datasets collected through high-frequency monitoring and routine sampling, we conducted rigorous comparative analyses using a comprehensive suite of statistical, machine-learning, and deep-learning models (Fig.1 and Table 1).

**Figure 1.**
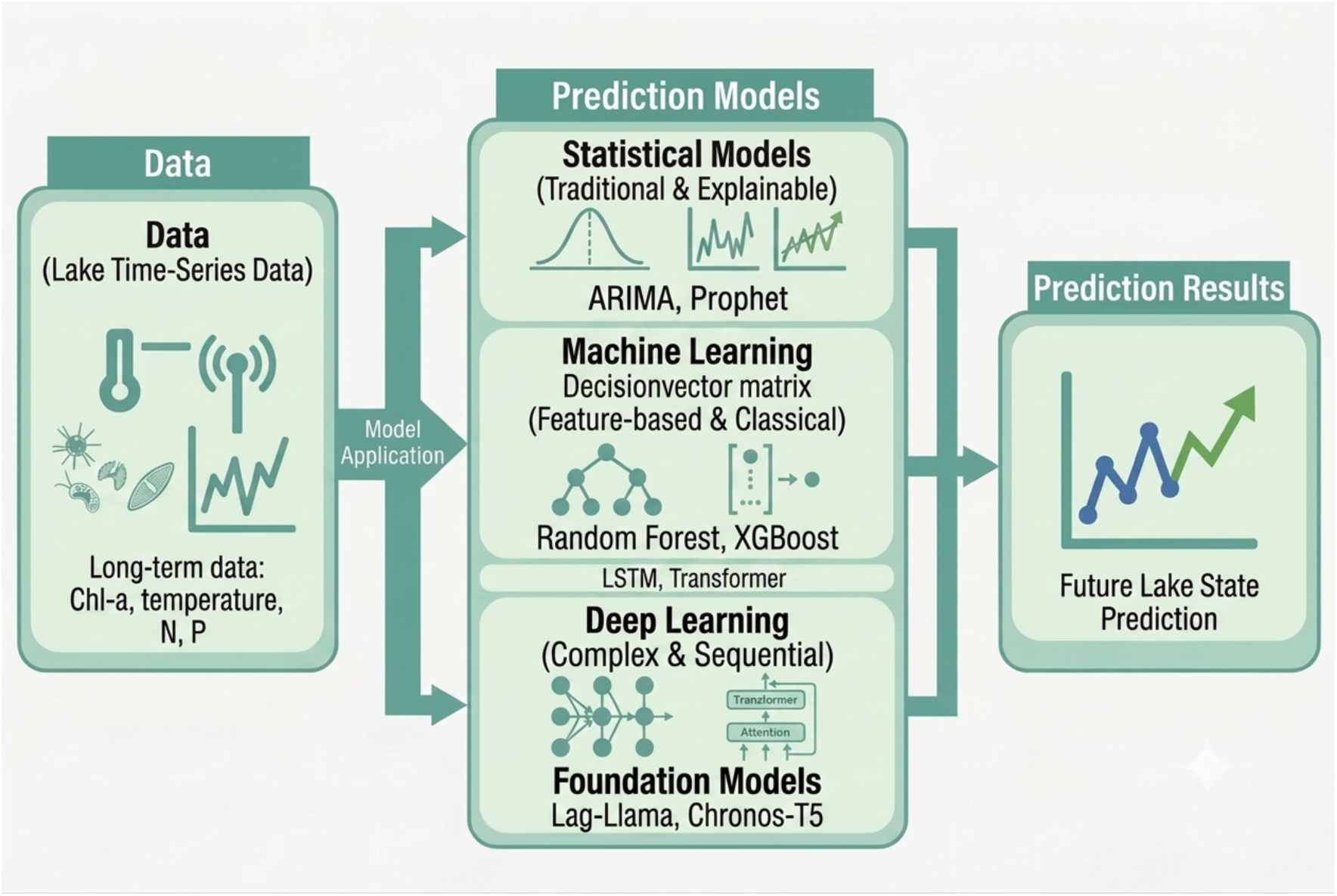
Conceptual framework of this study to forecasting the lake water quality. The framework illustrates the utilization of long-term time-series data (e.g., Chlorophyll-a [Chl-a], water temperature, nitrogen [N], and phosphorus [P]) as inputs for predicting future lake states. We compared 15 models (see Table 1) including statistical models (e.g., ARIMA, Prophet), machine learning models (e.g., Random Forest, XGBoost), deep learning (e.g., LSTM, Transformer), and foundation models (e.g., Lag-Llama and Chronos-T5).

**Table 1.**
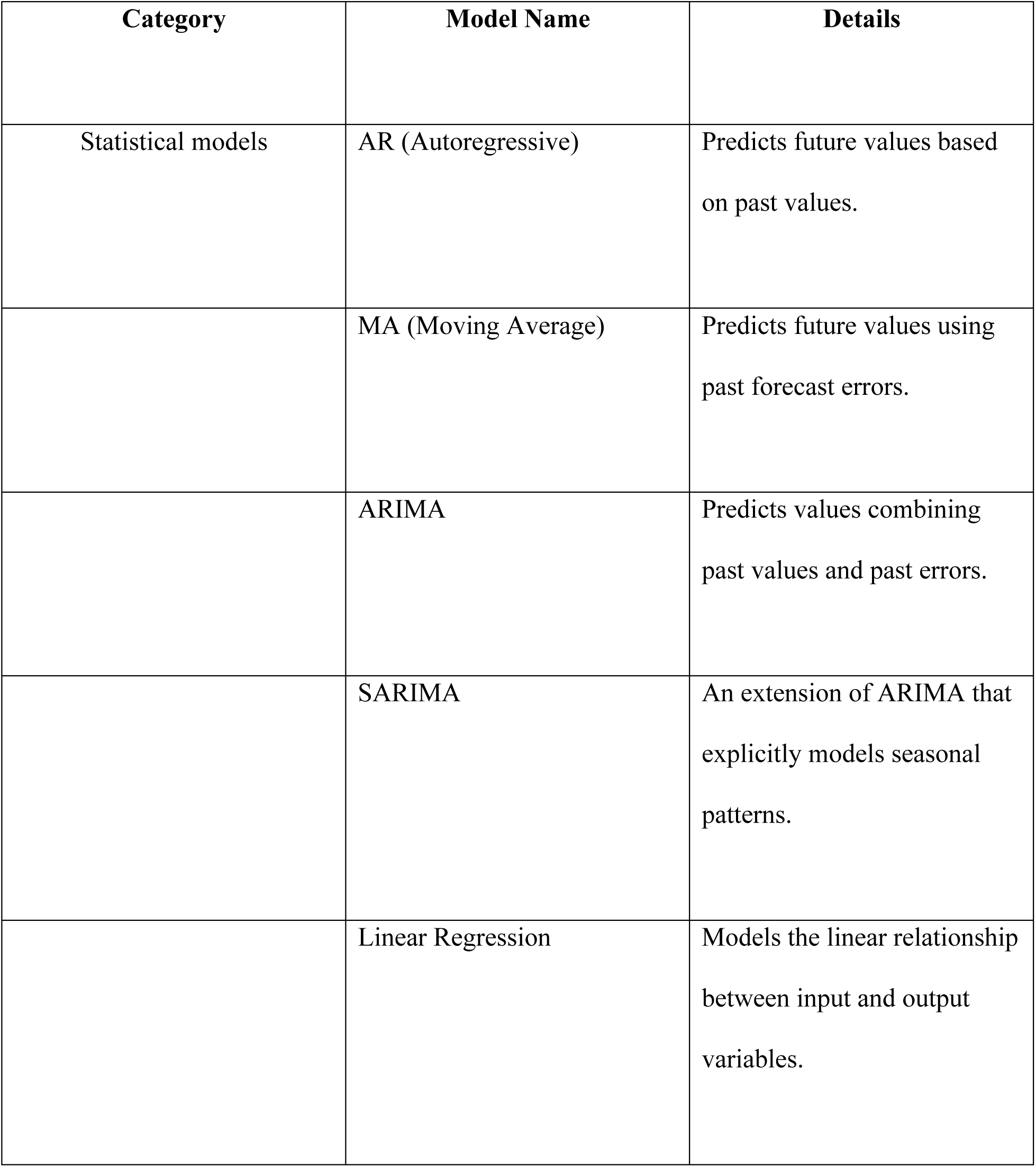
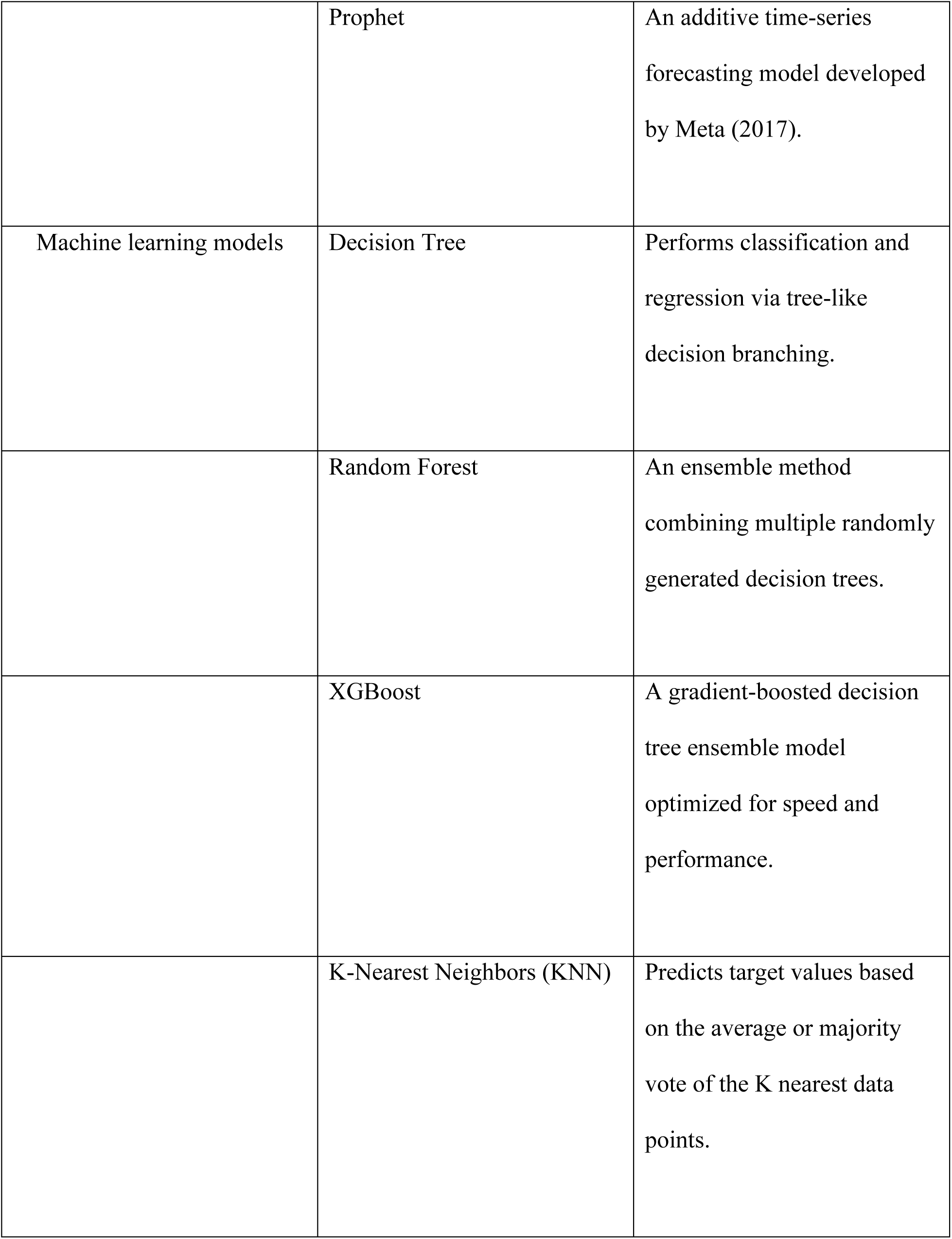
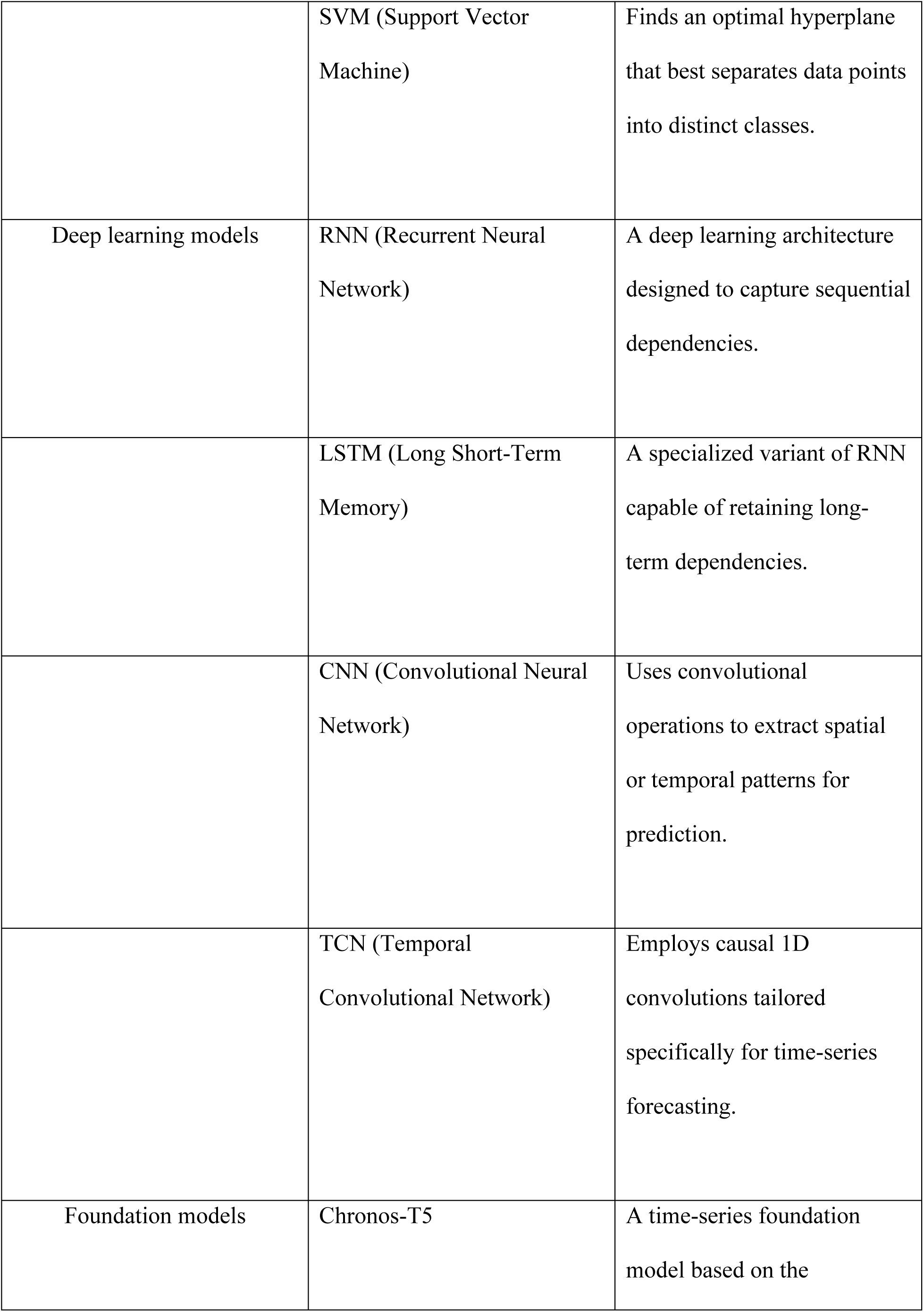
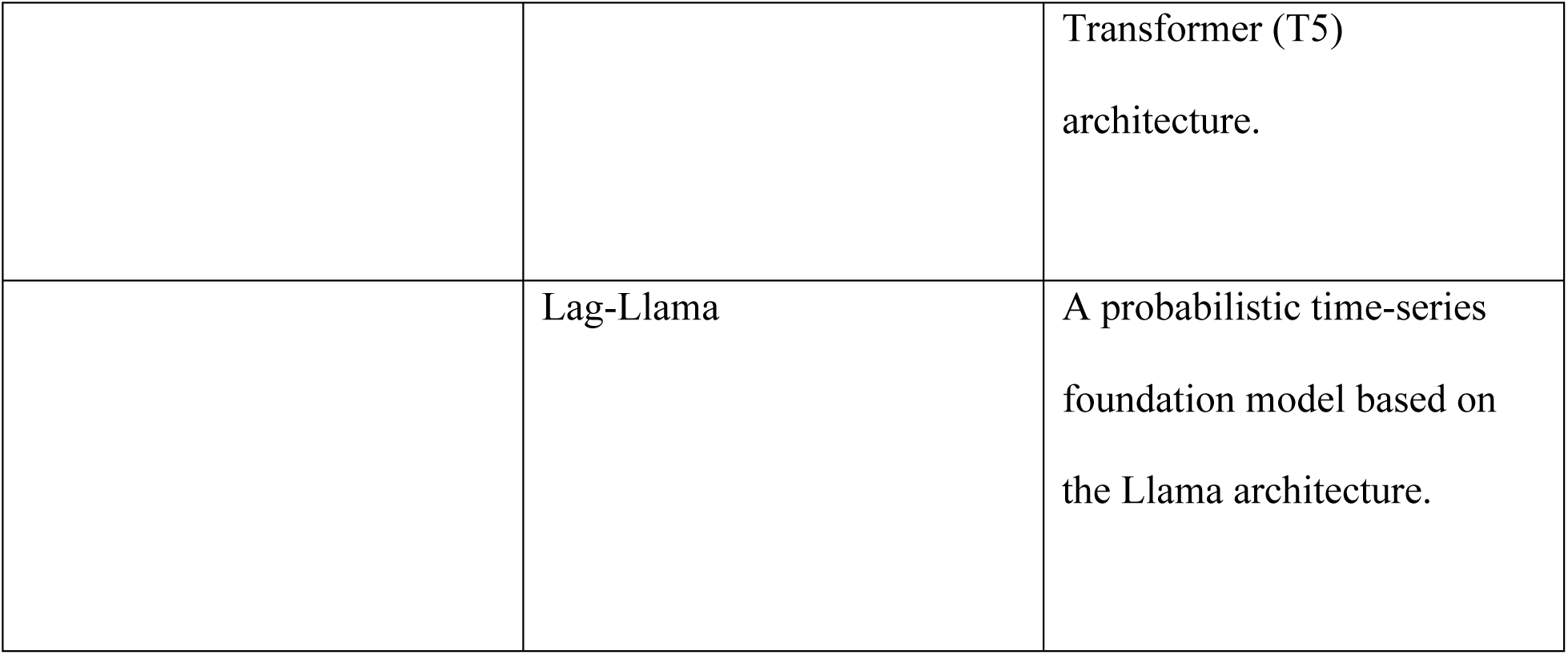
The 15 models spanning three distinct methodological categories using this study.

## Results

### Model Performance Comparison

Model performance comparisons across the 17 evaluated models reveal distinct capabilities and variable-dependent predictability across the six monitoring sites (Fig. 2 and 3). Notably, the fine-tuned Chronos-T5 model consistently dominated the overall performance, securing Rank 1 across all six sites (Fig. 2) and achieving exceptionally high predictive accuracy (R^2^ > 0.80–1.00, Fig. 3) for most parameters, including dissolved oxygen (DO), water temperature (WT), pH, and nutrients (NH4, NO3, and PO4). Interestingly, a sharp performance contrast exists within the foundation model category: while Chronos-T5 excels, Lag-Llama exhibits severe degradation (R^2^ ≈0), demonstrating that domain adaptation and underlying pre-training architectures critically dictate foundation model efficacy in aquatic time-series forecasting.

**Figure 2.**
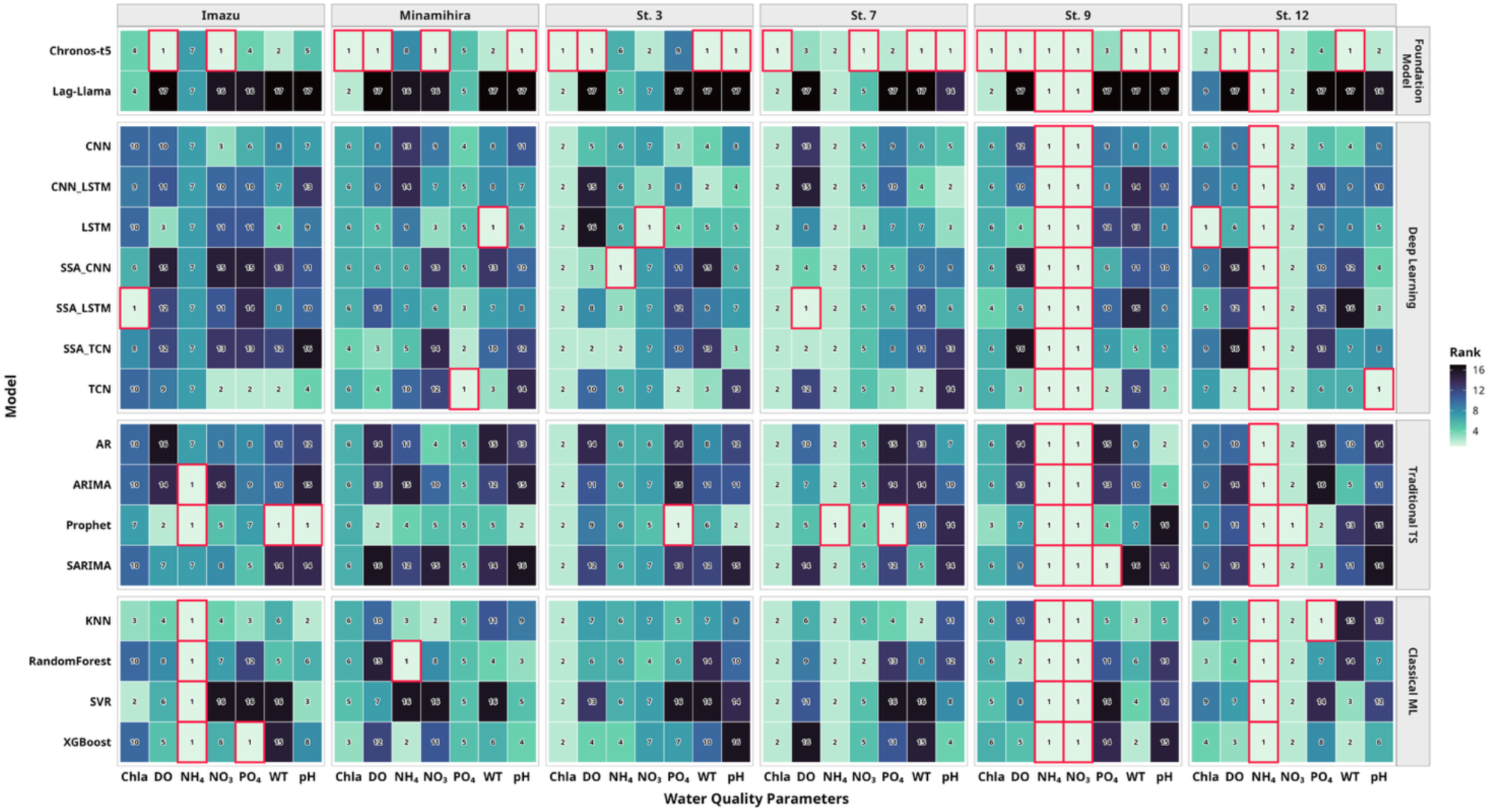
Heatmap for the rank of model performance (R²) for reproduction accuracy of 15 models for each site of the lake water quality indices in Lake Biwa and Lake Kasumigaura. Red squ are means the highest R² (i.e., top rank) among the methods.

**Figure 3.**
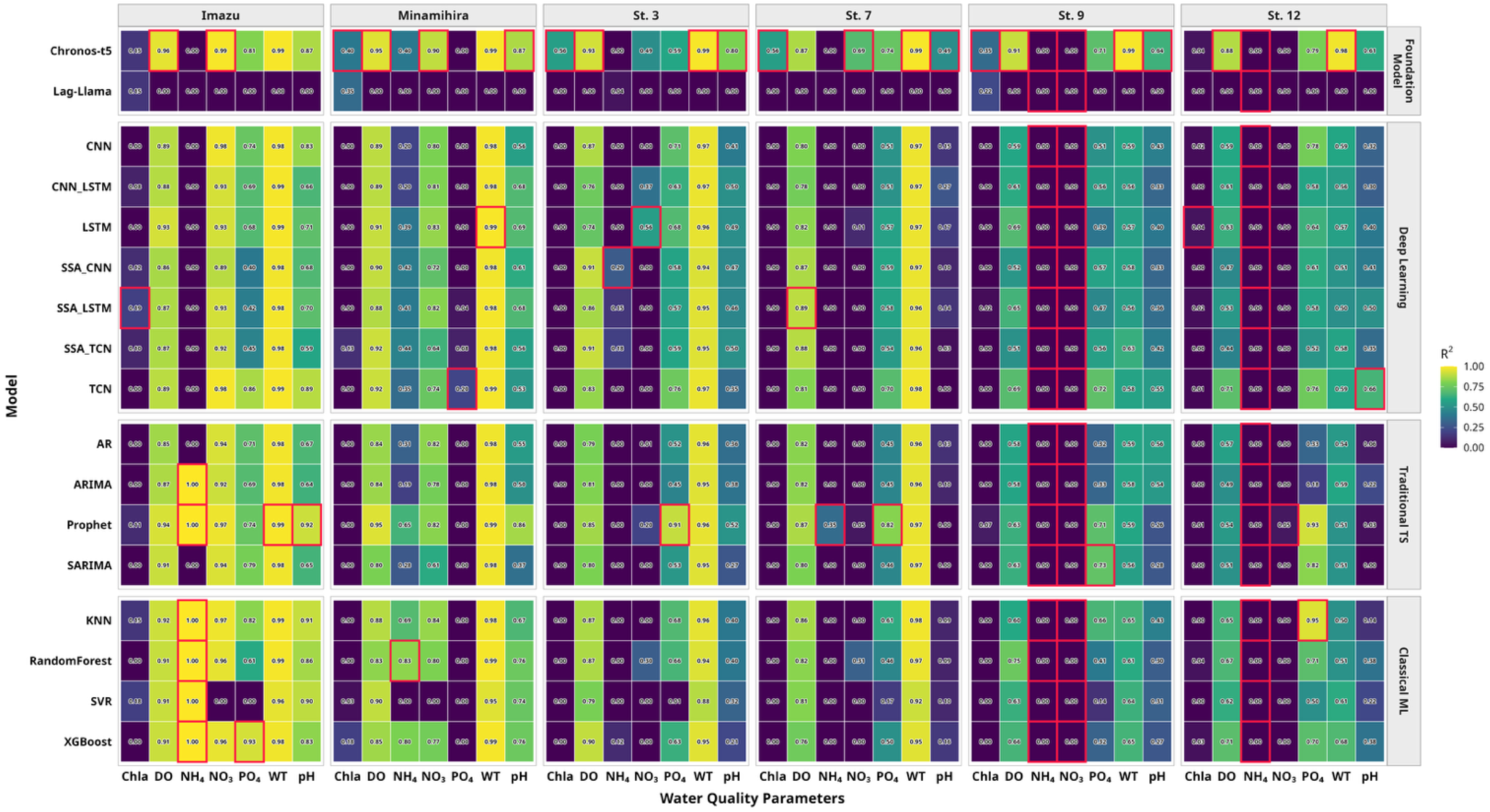
Heatmap for the model performance (R²) for reproduction accuracy of 15 models for each site of the lake water quality indices in Lake Biwa and Lake Kasumigaura. Blue square means the highest R²among the methods.

**Figure 4.**
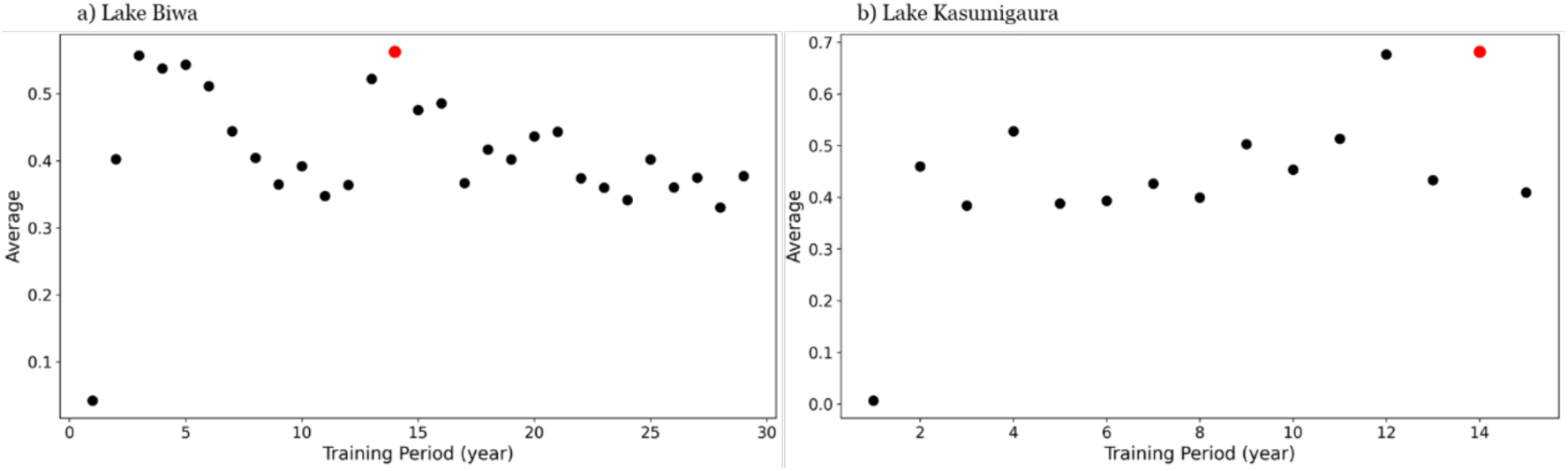
a) Average normalized recall accuracy by training period using Chronos-T5 for Lake Biwa and b) for Lake Kasumigaura. The red points indicate the maximum value within the training periods, representing the optimal training period as defined in this study.

Parameter-specific evaluations highlight substantial differences in modeling complexity across biological and chemical variables. Physical and stable chemical parameters, such as WT and DO, exhibit high baseline predictability across most deep learning, statistical models, and machine learning models. In contrast, chlorophyll-a (Chla) represents the most challenging variable, yielding uniformly low R^2^ values (R^2^ < 0.30) across all models, including Chronos-T5, due to its highly nonlinear, episodic biological dynamics driven by complex algal dynamics. For nutrient dynamics (NH4, NO3, and PO4), while Chronos-T5 maintains general superiority, classical machine learning models (e.g., KNN, XGBoost, Random Forest) and statistical methods (e.g., Prophet) occasionally achieve top ranks at specific localized sites (e.g., NH4 at Imazu or Minamihira). Overall, these findings indicate that while advanced foundation models such as Chronos-T5 provide superior generalizability across diverse water quality parameters, parameter-intrinsic complexity (such as in Chla) remains a primary bottleneck requiring specialized modeling strategies.

### Optimal Training Window for Chronos-T5

This pattern suggests that incorporating data spanning ecological regime shifts may introduce noise, thereby reducing model accuracy (Fig. 3). Therefore, selecting an optimal training window that balances sufficient data length with ecological stability is crucial. Future modeling efforts should consider regime dynamics to enhance their predictive performance. Specifically, the optimal training window was approximately 14 years for both lakes, with the model performance exhibiting a nonlinear relationship with the training data length. An initial local peak in performance occurred at 3–4 years, which may reflect short-term ecological consistency, followed by a decline as the training data incorporated the transitional periods. The global peak at 14 years likely represents a balance in which sufficient data are included to capture stable ecological patterns without the confounding influence of regime shifts. Extending training beyond 15 years degraded performance, likely reflecting ecological regime shifts that rendered older data less representative of current conditions. These findings emphasize the importance of carefully defining training periods in ecological modeling to avoid the detrimental effects of historical regime changes on predictive accuracy.

## Discussion

This study provides compelling evidence that large-scale time-series foundation models, particularly Chronos-T5, can facilitate transformative advances in ecological forecasting. Their superior performance is derived from their ability to leverage extensive pre-training across diverse domains, enabling robust long-term dependency modeling and generalization beyond the constraints of limited ecological data. This capacity to integrate heterogeneous datasets allows these models to capture complex temporal patterns and interactions that traditional models may overlook, thereby enhancing the depth and reliability of the ecological insights.

The superiority of Chronos-T5 over Lag-Llama underscores the profound influence of variations in model architecture, parameter scale, and training methodologies on the predictive performance of ecological time series models. Chronos-T5’s advanced architecture and larger parameter space allow it to model complex temporal dependencies and nonlinear interactions more effectively than Lag-Llama, which likely contributes to its enhanced accuracy and robustness in forecasting ecological dynamics. Furthermore, the substantial disparity in model size—Chronos-T5 at approximately 710MB versus Lag-Llama’s notably smaller 2.1MB—may play a critical role in this performance gap. The considerably larger size of Chronos-T5 suggests a greater capacity for learning intricate patterns and capturing subtle variations within the data, which smaller models, such as Lag-Llama, might be unable to represent fully due to their limited parameterization and representational power. This size difference not only reflects the complexity and depth of the underlying model but also implies potential trade-offs regarding computational requirements, memory usage, and deployment feasibility. Consequently, future research and model evaluations should incorporate systematic comparisons that consider model size alongside architectural design and training strategies. Such comprehensive analyses will enable a more nuanced understanding of how these factors collectively influence predictive accuracy, model generalizability, and computational efficiency, ultimately guiding the development of optimized models tailored to specific ecological forecasting tasks and resource constraints. As foundation models continue to increase in scale and complexity, their ability to capture intricate temporal dependencies and nonlinear ecological interactions is expected to improve substantially. This ongoing advancement, driven by innovations in algorithmic design and enhanced computational resources, points toward a future in which ecological forecasts become increasingly accurate, generalizable across diverse ecosystems, and robust to varying data conditions. Such improvements will be instrumental in informing more effective conservation strategies and resource management decisions, enabling stakeholders to respond proactively to environmental changes and biodiversity challenges. Moreover, the finding that smoothing through singular spectrum analysis (SSA) reduces forecasting accuracy emphasizes that ecological "noise" often contains vital biological signals essential for precise prediction. This challenges traditional preprocessing approaches that tend to suppress variability, underscoring the need to preserve subtle fluctuations that reflect real ecosystem dynamics and species interactions. Recognizing and appropriately modeling these meaningful variations is crucial for developing ecological models that authentically represent complex natural processes and improve long-term forecasting reliability.

The identification of a 14-year optimal training window aligns with the ecological theory of regime shifts, emphasizing that ecosystems undergo structural changes that limit the utility of long-term historical data. This insight has practical implications for model retraining and data selection strategies in ecological forecasting, encouraging adaptive approaches that balance historical depth and ecological relevance. Tailoring training windows to specific ecosystem dynamics may enhance model responsiveness to recent environmental changes, thereby improving forecast reliability and supporting more effective ecosystem management and policy formulation in the future.

Advances in GPU technology have mitigated traditional computational barriers, enabling researchers to deploy large foundation models on hardware configurations. This democratization of computational power has paved the way for the widespread adoption of these models in ecological research and management, facilitating real-time forecasting and informed decision making. Enhanced computational accessibility supports iterative model refinement and the integration of new data streams, fostering continuous improvements in ecological predictions and enabling timely responses to emerging environmental challenges. Additionally, these technological improvements reduce the costs and energy consumption associated with running complex models, thereby promoting more sustainable research practices. Consequently, smaller institutions and under-resourced research groups can contribute to and benefit from cutting-edge ecological modeling efforts, broadening the scope and diversity of scientific inquiry in this field. Given the variety of foundation models available, such as TimesFM, and the differences in model sizes even within the same architecture, it is essential to conduct comprehensive testing across multiple models and configurations to fully evaluate their performance and suitability for specific ecological applications in the future.

Future work should extend this framework across diverse ecosystems, taxa, and environmental conditions by integrating foundation models with mechanistic ecological understanding and climate projections to further enhance its predictive power and interpretability. Moreover, developing user-friendly pipelines and visualization tools is essential for translating complex model outputs into actionable management insights. Such interdisciplinary efforts will be pivotal in bridging the gap between advanced computational methods and practical ecosystem stewardship, ultimately harnessing the full potential of foundation models to address pressing environmental challenges and promote sustainable ecosystem resilience in a rapidly changing environment. Emphasizing capacity building and training for ecologists and resource managers in the use of these tools will be critical to ensure their effective implementation and adoption. Furthermore, fostering open data sharing and collaborative platforms can accelerate innovation and facilitate cross-regional comparisons, thereby strengthening global ecological monitoring and conservation strategies in the future.

The limitations of this study encompass several critical considerations that must be acknowledged to accurately interpret the findings. First, the scope of this study is confined to the selected context of the world, inherently limiting the generalizability of the findings to other geographical regions or specific subpopulations. Consequently, the outcomes may not fully capture the diverse cultural, economic, or social conditions prevalent in other areas, which could influence the applicability of the results. Second, potential biases may exist within the data collection and analysis methods employed, which could affect the objectivity and reliability of the findings. The diversity of analytical approaches used, while providing comprehensive insights, may also introduce variability in the interpretation and comparability of results across different studies or contexts. Third, the study may not have accounted for all relevant variables or confounding factors that could influence the observed outcomes, thereby limiting the depth of interpretation and the robustness of the conclusions drawn. Additionally, temporal constraints pose a significant limitation; data were gathered within a specific time frame, restricting the ability to generalize trends over extended periods or to capture dynamic changes that may occur in evolving contexts. Furthermore, reliance on secondary data sources or self-reported information introduces potential inaccuracies, inconsistencies, or reporting biases that could compromise data reliability and validity. Finally, the absence of longitudinal data limits the study’s capacity to infer causality or examine long-term effects, thereby reducing the strength of any causal claims and hindering the understanding of temporal relationships within the studied phenomena.

In conclusion, time-series foundation models represent a transformative advancement in ecological forecasting, delivering exceptional accuracy and robustness in predicting complex and inherently noisy biological dynamics. The Transformer-based Chronos-T5 model exemplifies this progression by harnessing the power of large-scale pre-training to effectively address the traditional challenges associated with limited and heterogeneous ecological data. This approach establishes a scalable, data-driven framework that is well-suited for the comprehensive management of aquatic ecosystems. By identifying and implementing optimal training strategies, this study enhances the predictive capabilities of foundation models and demonstrates their practical feasibility in real-world ecological contexts. Such advancements provide a critical foundation for integrating these models into operational frameworks, including early warning systems capable of timely detection of ecological disturbances and adaptive resource management protocols that respond dynamically to environmental changes. By facilitating improved forecasting accuracy and operational readiness on a global scale, the Chronos-T5 model and similar time-series foundation models hold significant promise for advancing sustainable ecosystem management. This study lays the groundwork for the broader adoption and continued development of foundation models, ultimately contributing to more resilient and well-informed stewardship of aquatic environments worldwide.

## Methods

### Study Sites and Data Collection

This study focused on phytoplankton in domestic lakes and selected study sites that satisfy the criteria of having monthly continuous chlorophyll-a concentration data— used as an index of phytoplankton biomass—for a period of 10 years or more as time-series data. Data on water quality parameters from multiple sites in two lakes, Lake Biwa and Lake Kasumigaura, that met these conditions were analyzed.

For Lake Biwa, among the monitoring sites in the Environmental Survey Information Database of the Lake Biwa Environmental Research Center (https://www.lberi.jp/investigate/water), two sites where continuous monthly surveys were conducted are Imazu-oki and Minami-Hira-oki (Fig. S1). Regarding the water quality parameters, database records were retrieved for chlorophyll-a concentration (Chl.a: µg L⁻¹) and factors identified in Section 1.2 that can influence seasonal fluctuations in phytoplankton biomass and are available as time-series data: water temperature (WT: °C), dissolved oxygen concentration (DO: mg L⁻¹), pH, phosphate phosphorus concentration (PO₄³⁻-P: µg L⁻¹), ammonium nitrogen concentration (NH₄⁺-N: µg L⁻¹), and nitrate nitrogen concentration (NO₃⁻-N: µg L⁻¹). Data collected at a depth of 5 m were used, covering a 30-year period of monthly data from January 1994 to December 2023.

For Lake Kasumigaura, the data from the Global Environmental Database (GED) of the National Institute for Environmental Studies (https://db.cger.nies.go.jp/gem/inter/GEMS/database/kasumi/index.html) were used. Four survey sites where water quality was monitored continuously on a monthly basis— St. 3, 7, 9, and 12 (Fig. S1)—were selected. Database records were retrieved for chlorophyll-a concentration (Chl.a: µg L⁻¹), water depth (m), and factors identified in Section 1.2 that can influence seasonal fluctuations in phytoplankton biomass and are available as time-series data: water temperature (WT: °C), dissolved oxygen concentration (DO: mg L⁻¹), pH, phosphate phosphorus concentration (PO₄³⁻-P: µg L⁻¹), ammonium nitrogen concentration (NH₄⁺-N: µg L⁻¹), and nitrate nitrogen concentration (NO₃⁻-N: µg L⁻¹). Analysis was conducted using 16 years of monthly data from January 1997 to December 2012.

### Model Selection and Implementation

We implemented two state-of-the-art time-series foundation models to forecast ecological variables: Chronos-T5, which utilizes a Transformer-based architecture capable of capturing complex temporal dependencies,^16^ and Lag-Llama, a Llama-based probabilistic forecasting model designed to generate uncertainty-aware predictions^15^. These advanced foundation models were benchmarked against a comprehensive suite of 15 alternative models spanning three distinct methodological categories (Table 1). The statistical model group included classical time-series approaches, such as autoregressive (AR), autoregressive integrated moving average (ARIMA), seasonal ARIMA (SARIMA), and Prophet models, which are widely used for their interpretability and efficiency in capturing linear and seasonal trends. The machine learning group comprised algorithms such as random forest, XGBoost, K-nearest neighbors (KNN), and support vector regression (SVR), which offer nonlinear modeling capabilities but often require careful feature engineering and data preprocessing. Finally, the deep learning group incorporated advanced architectures, such as long short-term memory (LSTM) networks, convolutional neural networks (CNNs), temporal convolutional networks (TCNs), and hybrid models combining these with singular spectrum analysis (SSA) for noise reduction. This diverse model selection enabled a thorough comparative evaluation of predictive performance across different algorithmic paradigms, highlighting the strengths and limitations of each approach in handling complex, noisy ecological time series data.

Nutrient concentrations below the detection limit were set to zero to maintain analytical consistency and ensure uniformity across the datasets. The trend component extracted from the time series revealed gradual, long-term changes that likely reflect underlying shifts in environmental conditions such as climate variations, land use changes, and anthropogenic impacts. These trends provide valuable insights into the persistent directional movements of water quality parameters over a multi-decadal study period. Seasonal patterns identified through decomposition were consistent with established cyclical environmental variations, including predictable fluctuations in temperature, light availability and nutrient cycling. These seasonal components exhibited regular periodicity, aligning with annual climatic and ecological rhythms that influence the biological productivity and chemical dynamics of lakes. The residual component captured irregularities and short-term anomalies that were not accounted for by the trend or seasonal factors in the model. These residuals likely represent episodic events, such as storm runoff, algal blooms, or measurement noise, highlighting the presence of stochastic variability and transient disturbances in the aquatic ecosystems. Understanding these residual fluctuations is essential for interpreting deviations from expected seasonal and long-term patterns.

Chronos-T5 was fine-tuned with a 12-month forecast horizon using a batch size of 2 and a learning rate of 0.001, leveraging the pre-trained amazon/chronos-T5-large model. Lag-Llama generated probabilistic forecasts by sampling 100 trajectories and averaging the results. For Choronos-T5, the model parameters are set as Table 2. Statistical and machine learning models incorporated engineered features, such as lagged variables, moving averages, and monthly dummies, with appropriate normalization applied where necessary.

**Table 2.**
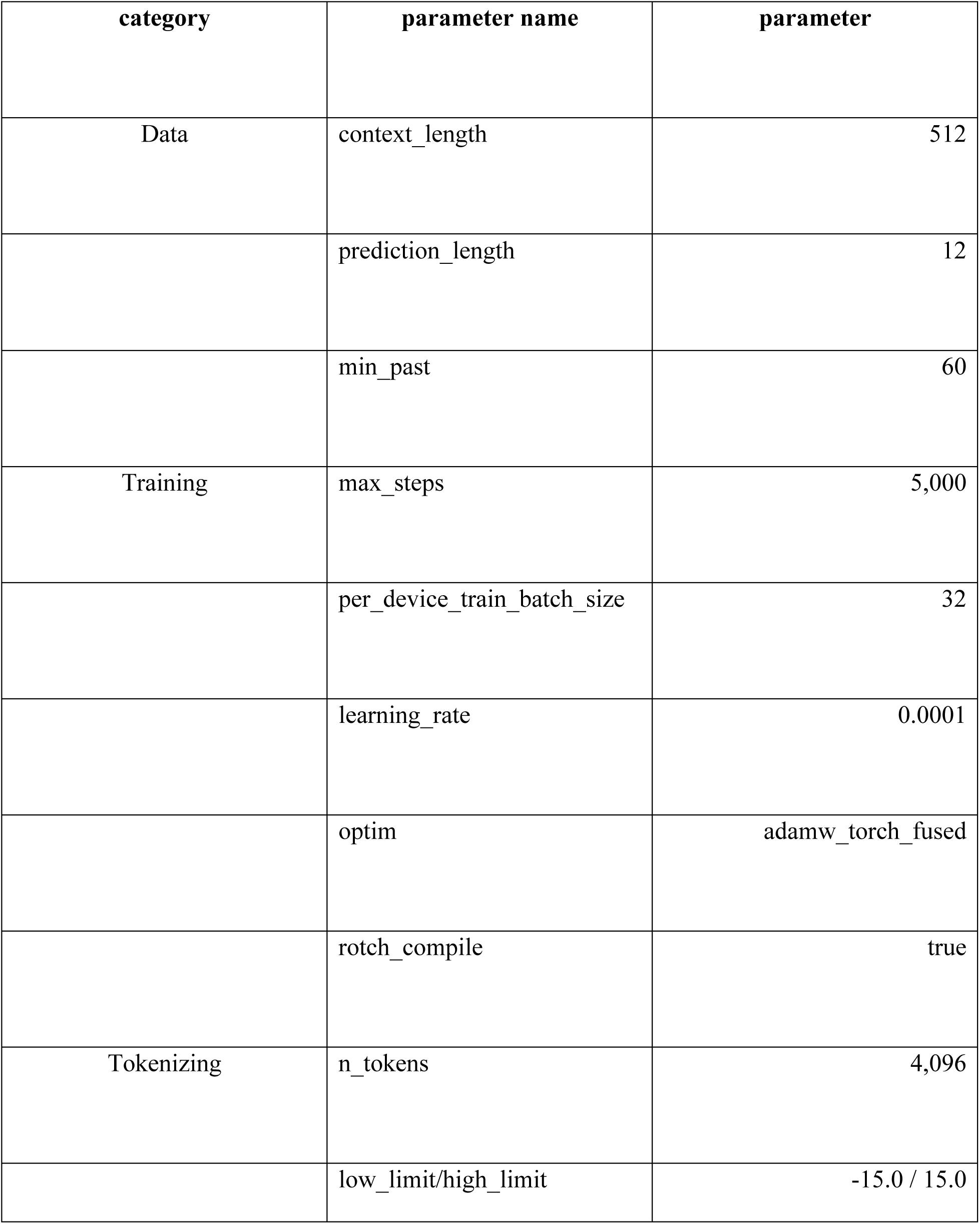
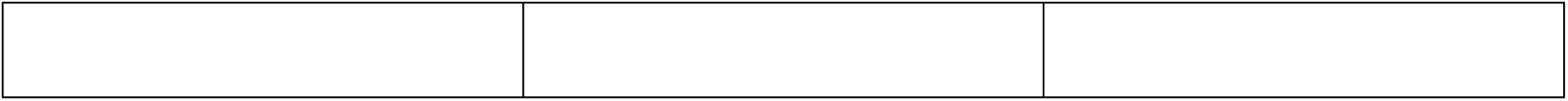
Details on training and tokenizer Parameters for Chronos-T5.

| category | parameter name | parameter |
| --- | --- | --- |
| Data | context_length | 512 |
|  | prediction_length | 12 |
|  | min_past | 60 |
| Training | max_steps | 5,000 |
|  | per_device_train_batch_size | 32 |
|  | learning_rate | 0.0001 |
|  | optim | adamw_torch_fused |
|  | rotch_compile | true |
| Tokenizing | n_tokens | 4,096 |
|  | low_limit/high_limit | -15.0 / 15.0 |

#### Training and Validation

Training windows varied extensively, ranging from 1 to 29 years for Lake Biwa and 1 to 15 years for Lake Kasumigaura, allowing for a comprehensive exploration of temporal scales in model training. Subsequent validation was rigorously performed on holdout years, specifically 2023 for Lake Biwa and 2012 for Lake Kasumigaura, to ensure that the predictive capabilities of the models were tested on independent data. Model performance was quantitatively evaluated using the coefficient of determination (R^2^), with any negative values clipped to zero to prevent misleading interpretations and uphold the reliability and clarity of the evaluation metric. The identification of optimal training periods was guided by peak normalized values, which served as a robust indicator balancing the need for ample data volume against the necessity of maintaining stable environmental regimes throughout the training window. This approach mitigates the risk of overfitting by avoiding anomalous or abrupt regime shifts that could distort model generalizability. Moreover, the selected training windows reflect a strategic compromise between maximizing the amount of available data and preserving the ecological and environmental consistency critical for reliable model performance. This careful calibration enhances the robustness and applicability of the models across varying temporal contexts, thereby supporting more accurate and stable predictions.

## Data Availability

Previously published data were used for this work as described in Data collection of the Method.

## Code Availability

Custom scripts used for data processing and model evaluation are available from the corresponding author upon reasonable requests.

## Acknowledgements

This research was funded by the Japan Science and Technology Agency (JST) under the CREST grant number JPMJCR24J2 and the Environment Research and Technology Development Fund of the ERCA (JPMEERF 26S21300), funded by the Ministry of the Environment.

## Author Contributions

A.M. and H.D. contributed equally to this study. A.M. conducted the data analyses. Both authors participated in discussions regarding the results, developed the conceptual framework, and contributed to preparing the final manuscript.

## Disclosure of AI usage

The authors used AI tools, including paperpal.com and Google Gemini, to assist with polishing language, condensing text, Figure 1 illustration and improving clarity during the preparation of this manuscript. All outputs were reviewed, adapted, and integrated into the final work by the authors, who took full responsibility for all the content presented herein. Note: Disclosure of AI-assisted copy editing for grammar, style, readability, and formatting is not required.

## Supplemental Materials

**Figure S1.**
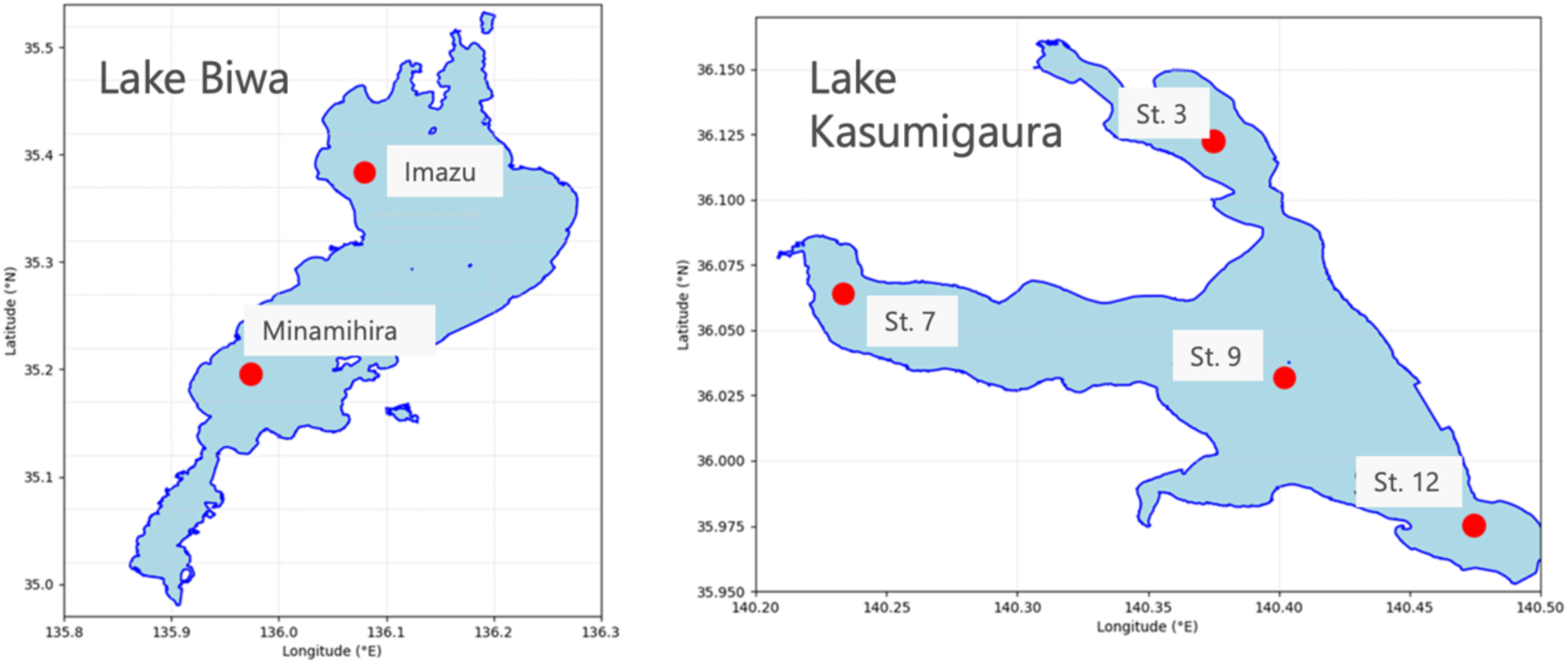
Study sites for the data used in this study. Data from two sites in Lake Biwa, St. Imazu and Minamihira (left) and four sites in Lake Kasumigaura, St. 3, 7, 9, and 12 (right).

